# Complex genetic architecture underlying the plasticity of tobacco leaf width provides insight into across-environment genomic prediction

**DOI:** 10.1101/2024.05.05.592603

**Authors:** Li Liu, Yu Han, Wei Liu, Yanjun Zan, Dong Wang, Yan Ji, Yiwen Sun, Xin Liu, Ran Hao, Wenjia Zhang, Linjie Guo, Jiarui Zhao, Zhimei Song, Dan Liu, Aiguo Yang, Caineng Zhao, Haizhou Hu, Lirui Cheng, Huan Si

## Abstract

Phenotypic plasticity refers to the ability of a given genotype to produce multiple phenotypes in response to changing environmental conditions. Understanding the genetic basis of phenotypic plasticity and developing predictive models for agronomic traits are crucial for future agricultural adaptation to climate change. In this study, we investigated the genetic basis of leaf width mean (LWm) and plasticity (LWp) in a tobacco multiparent advanced generation inter-cross (MAGIC) population which consisting of 594 individuals. We identified 14 quantitative trait loci (QTLs) significantly associated with LWm, 43 with LWp. Our findings suggest that dynamic changes in QTL effects across environments, along with polygenic effects, may underlie the genetic basis of leaf width plasticity. Among them, *qLW14* was narrowed down to a 3 Mb structural variation region. When this fragment was deleted in tobacco, plants exhibited increased leaf width, but only under specific environmental conditions. This finding suggests that the key gene within *qLW14* may act as a negative regulator of leaf width through interactions with specific environmental factors. By integrating genetic diversity, environments variation, and their interactions into a GEAI model, we were able to build a framework for cross-environment prediction, improving prediction accuracy by 8.7% when compared to traditional model. Overall, this study highlights the complex genetic basis underlying LWp, involving multiple alleles and genotype-environment interactions. These findings provide valuable insights into the role of phenotypic plasticity in plant adaptation to environmental changes.

## 1. Introduction

Phenotypic plasticity refers to the phenomenon where individuals of the same genotype exhibit different phenotypes under different environmental conditions (Bradshaw, 1965; Pigliucci, 2005), or the ability of organisms to produce different phenotypes in response to varying environmental factors (Kliman, 2016; Nicotra et al., 2010). In organisms, phenotypic plasticity is a trait and possibly an important strategy to quickly adjust phenotypes according to environments, which potentially have effects on evolution (Kusmec et al., 2018; Sommer, 2020). As sessile organisms, plants have evolved unique mechanisms to cope with rapid climate changes, including physiological, metabolic, growth, and developmental adjustments (Brooker et al., 2022; Des Marais et al., 2013). To understand the mechanism of plastic responses is important because agricultural systems will face immense pressure from rapid population growth and climate change in the future.

Previous studies on model organisms, such as yeast, and major crops like rice, maize, and cotton have demonstrated that phenotypic plasticity is widespread and under complex genetic regulation (Alvarez Prado et al., 2014; Kikuchi et al., 2017; Kusmec et al., 2017; Pigliucci, 2005; Zan and Carlborg, 2020). However, fully leveraging phenotypic plasticity requires a comprehensive understanding of its genetic architecture. Three genetic models have been proposed to explain the regulation of phenotypic plasticity, including overdominance model (Gillespie and Turelli, 1989), allelic sensitivity model (Via, 1993) and regulatory gene model (Scheiner and Lyman, 1989; Schneider et al., 2020). The overdominance model suggests that heterozygosity at genes causes plastic responses. However, both the allelic sensitivity model and the regulatory gene model describe genotype-environment interaction (G×E) as the variation in how different genotypes respond to environmental changes. The key distinction lies in their proposed mechanisms: the allelic sensitivity model attributes plastic responses to environmentally sensitive alleles of genes that influence mean phenotypes, whereas the regulatory gene model suggests that plasticity arises from the regulation of mean phenotype genes by other genes that integrate environmental stimuli.

When comparing the genetic structures underlying phenotypic means and plasticity, some studies have demonstrated that they are neither entirely unified nor entirely independent, but rather exhibit partial overlap (Alvarez Prado et al., 2014; Guo et al., 2020; Jin et al., 2023). For example, a study on maize grain size and weight plasticity revealed that 30% of the QTLs associated with plasticity were co-localized with those governing phenotypic means, indicating a shared genetic basis (Li et al., 2019). However, in another research on 23 maize traits, only 6% of the QTLs associated with plasticity also co-localized with phenotypic mean loci, which showed that the candidate genes associated with mean phenotypes and their plasticity are structurally and functionally distinct across all 23 phenotypes (Kusmec et al., 2017). To explore the genetic mechanisms underlying phenotypic plasticity have significant implications for plant breeding.

In recent years, the combination of genome-wide association study (GWAS) and genome selection (GS) has been widely reported to play a significant role in accelerating the breeding processes, by integrating the QTLs from GWAS into GS programs to enhance the prediction accuracy of phenotypes (Desta and Ortiz, 2014; Huang et al., 2016, 2023; Resende et al., 2012; Spindel et al., 2016; Tong et al., 2020; Wang et al., 2015; Si et al., 2025). However, the prediction performance of GS model often varies significantly across different environments, primarily due to the lack of consideration for G×E. Therefore, uncovering the genetic basis of phenotypic plasticity and integrating plasticity-associated QTLs into GS models is of great significance for enhancing the accuracy of phenotypic prediction in heterogeneous environments.

Tobacco (*Nicotiana tabacum* L.) originated from interspecific hybridization between *Nicotiana sylvestris* and *Nicotiana tomentosiformis*. As an allotetraploid species, tobacco exhibits high diversity and plasticity in leaf morphology, plant architecture, and metabolic traits (Honarnejad and Shoai-Deylami, 2004; Legg and Collins, 1975; White et al., 1979), making it an important model organism for research in plant genetics and developmental biology. Over the past two decades, advancements in high-throughput sequencing technologies and the decreasing cost of sequencing have significantly facilitated research on the genetic mechanisms underlying key traits in plants and animals, including tobacco (Edwards et al., 2017; Khan et al., 2019; Resende et al., 2012; Wang et al., 2022). Leaves are the primary harvestable organ of tobacco and play a crucial role in its growth, development, and agricultural production. Recently, GWAS have been widely applied to investigate the genetic basis of leaf traits in tobacco, leading to the identification of numerous QTLs associated with leaf number, width, and length (Ahmed et al., 2019; Liu et al., 2022b; Safdar et al., 2020; Zuo et al., 2022), which provided valuable resources for revealing the genetic basis of plant growth and development. However, the majority of these genetic studies have focused on mean values of phenotypes, while the genetic basis of plasticity of traits remains largely unexplored.

In this study, we conducted a GWAS in a tobacco MAGIC population consisting of 594 individuals and identified a total of 45 QTLs for LWm, LWp and. Our findings suggest that dynamic changes in major QTL effects across environments, along with polygenic effects, may underlie the genetic basis of leaf width plasticity. Further analysis identified the candidate region of *qLW14* associated with leaf width plasticity Further analysis identified the candidate region of *qLW14* associated with leaf width plasticity and refined it to a ∼3 Mb structural variation located at the distal end of chromosome 14.. By integrating genetic diversity, environmental variation, and their interactions, we developed a framework for cross-environment prediction, improving prediction accuracy by 8.7% compared to traditional models. Overall, this study elucidates the complex genetic architecture underlying both leaf width mean and plasticity, involving multiple alleles and G×E. These findings offer valuable insights into the role of phenotypic plasticity in plant adaptation to environmental changes and provide a theoretical foundation for its application in crop breeding.

## 2. Materials and methods

### 2.1 Plant materials and growth conditions

A total of eight different tobacco varieties were utilized as parents to develop the MAGIC population. These included *Beinhart1000-1* and *Florida301*, which is cigar tobacco varieties; *Basma* and *Samsun*, belonging to oriental tobacco; *Xiaohuaqing* (XHQ) and *Tangpeng* (TP) from sun-cured tobacco; *Vam*, a variety of Virginia tobacco; and finally, the flue-cured tobacco variety *Honghuadajinyuan* (HD). Initially, these eight parents were crossed in pairs to produce four two-way crosses. The offspring from these crosses then underwent incomplete diallel crossing to generate four-way crosses, which were further crossed to produce eight-way crosses. These eight-way crosses were then subjected to random mating, yielding a total of 320 eight-way crossbreds. From these 320 eight-way crossbreds, three individual plants were selected per crossbred, and then propagated for five generations through the single seed descent (SSD) method, resulting in the establishment of over 800 eight-way tobacco MAGIC inbred lines.

In 2019 and 2020, the MAGIC population, along with the eight parents, was planted separately in Zhucheng, Shandong Province, China, and Guiyang, Hunan Province, China, respectively. A randomized complete block design was employed with two replicates. Each replicate consisted of teo rows of 10 plants per line, with a row length of 10 m, a row spacing of 1.2 m, and a plant spacing of 0.5 m.

### 2.2 Phenotype and Genotype

During each experiment, five randomly selected plants were selected for phenotypic evaluation, and the average value was analyzed. Leaf width was defined by measuring the width of the blade perpendicular to the main vein for the middle leaf. In addition to LWm, we assessed LWp using SP and OP (Methods 2.5), which quantify the variation in LW across different environments.

Genotyping was conducted using a tobacco SNP chip containing 432,362 markers from the Zhengzhou Tobacco Research Institute of the China Tobacco Corporation. Detailed information regarding genotyping and SNP calling procedures can be found in a previous study (Yuan et al., 2023).

### 2.3 Calculation of narrow-sense heritability and genetic correlation

We used a mixed linear model to estimate the narrow-sense heritability:

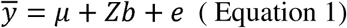

Among them, *y̅* represents LWm, with a total of 594 rows of data. µ is the intercept, which represents the population mean. *Z* is the design matrix, satisfying *ZZ^T^* = *G*, where *G* is the genetic relationship matrix estimated by GCTA (Yang et al., 2011). Therefore, *b* follows a normal distribution 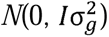, and *e* is the residual, which follows 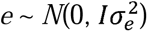. The narrow-sense heritability (*h^2^*) is calculated by the interclass correlation, with the formula 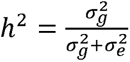. All analysis were performed in GCTA. Furthermore, we used multivariate mixed linear models in ASReml to estimate pairwise genetic correlations between LWm measured in the four environments.

### 2.4 Phenotypic Variance Analysis and Estimation of BLUP

To quantify the contribution of each factor to phenotypic variation, we conducted an analysis of variance (ANOVA) based on a linear model. The model included genotype (id), environment (loc), and their interaction (id * loc) as explanatory variables:

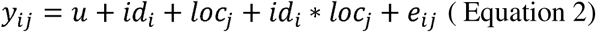

*y_ij_* (*i*= 1…n, n=560, number of individuals; *j*=1…m, m=4,number of sites) represents the leaf width measured for each individual across four environments; u represents the population mean; The *id_i_* term denotes the genotype of each individual and was treated as a categorical variable to capture genetic differences among lines; The *loc_j_* term represents the environmental factor, and the interaction term *id_i_ * lac_j_* captures genotype-by-environment interactions (G × E); The *e* term accounts for residual error. The proportion of total phenotypic variance explained by each component was estimated using the sum of squares (SS) obtained from the ANOVA.

We used the following model to estimate the BLUP of leaf width:

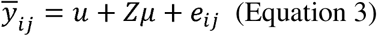

where *y̅_ij_* represents the leaf width measurements of 594 individuals across four environments; u represents the population mean; *Z* is the design matrix linking the BLUPs estimates with the observed phenotypic values; µ is a vector of random effects representing the BLUP estimates of 594 individuals, assumed to follow a normal distribution 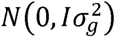; and *e_ij_* represents the residual error.

### 2.5 Quantification of plasticity measurements

Since all the MAGIC lines were phenotyped for LW across four environments, we quantified and studied the genetics of LW plasticity in response to environmental variation. Here, the phenotypic plasticity was classified into two categories. (1) overall plasticity, which describes plasticity across all the studied environments, and (2) the second category-specific plasticity is more unique to certain pairs of sites, which only captures the plasticity across two environments. The motivation underlying such classification is that some individuals are more robust across most of the studied sites except only one or a few sites, while other individuals are plastic among most of the sites. Site-specific plasticity was quantified using pairwise differences in phenotypic values between two environments. In addition, two additional approaches were used to quantify the overall plasticity. First, the across environment variance (VarR) (Vanous et al., 2019) of the rank transformed phenotype was used to account for the mean difference. The fourth score of the overall plasticity (FWR) (Finlay and Wilkinson, 1963; Lian and de Los Campos, 2016) applies Finlay-Wilkinson Regression to partition the phenotype into two components, one that constant across environments and another that responds dynamically to environmental changes. Using the linear mixed model, the phenotype of each line is partitioned into these two components and the plasticity component is used as a measurement of plasticity. In total, the described approaches resulted in two measurements of phenotypic plasticity, abbreviated as SP, OP.

### 2.6 Genome-wide association analysis for LW mean/BLUP and plasticity measurements

Genome-wide association analysis was conducted using a linear mixed model implemented in GCTA. We applied a significance threshold of p < 0.05/Me for SNPs, where Me represents the number of independent SNPs estimated following methods described by Li et al. (Li et al., 2012). Effect sizes of identified quantitative trait loci (QTL) were extracted from the GCTA output. The effect sizes for each SNP were estimated using the previously mentioned mixed linear model.

### 2.7 Predicting the site-specific performance of the LW

We fitted the following models to predict leaf width phenotypes:

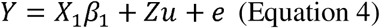

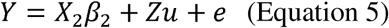

Here, *Y* is a phenotypic value vector of size 560 ×4, representing the leaf width measurements of 560 individuals across four environments. The term *X*_1_*β*_1_ represents the fixed effects component, where *X*_1_ is the design matrix for fixed effects, including a column of 1 to represent the overall mean and three additional columns to represent the environmental effects of the remaining three sites relative to a reference site. The vector *β*_1_ contains the corresponding fixed effect coefficients, indicating the contribution of each fixed effect to the phenotypic values. The term *Zu* represents the random effects component, where *Z* is the design matrix satisfying the condition *ZZ*′ = *G* ⊗ I, with G being the genomic relationship matrix (GRM)(Yang et al., 2011) that captures the genetic similarity among individuals and I being the identity matrix. The vector *u* represents the random effects, specifically the breeding values of the individuals, reflecting their genetic potential contribution to the phenotypic values. The term e represents the randomly distributed residuals.

We define equation (4) as the GBLUP model, while equation (5) is defined as the GEAI model. Compared to *X*_1_ in equation (4), *X*_2_ in equation (5) is an extended design matrix that incorporates QTL genotypes and QTL-by-environment interaction terms in addition to the columns in *X*_1_. This extension allows the GEAI model to simultaneously capture systematic effects, genetic effects, and genotype-by-environment interactions, thereby providing more accurate predictions, particularly in cross-environment and novel environment predictions. These models were fitted using the rrBLUP (Endelman, 2011) package in R (https://www.R-project.org/).

We employed two methods to predict leaf width phenotypes. In the first method, for each of the four environments, 80% of the individuals were randomly selected as the training set, while the remaining 20% were used as the test set, with their phenotypic values masked as *NA*. The coefficient of determination (*r*^2^) between the predicted and measured phenotypes was used to evaluate prediction accuracy within each environment. In the second method, all individuals from any three of the four environments were used as the training set, while all individuals from the remaining environment were used as the test set, with their phenotypic values masked as *NA*. The coefficient of determination (*r*^2^) between the measured and predicted phenotypes within each site was used to assess prediction accuracy.

## 3. Results

### 3.1 Tobacco leaf width mean variation within each of the four environments

To investigate the genetic basis of leaf width mean (LWm) and plasticity (LWp), we analyzed 594 tobacco lines derived from a tobacco MAGIC population. Genotypes were performed using the 430K tobacco SNP array, which includes 432,362 markers, Leaf width was measured under four different environments in Guiyang (25°27’N, 112°13’E) and Zhucheng (35°59′N, 119°24′E) during 2019 and 2020 (Figure 1A, Table S1). The studied environments were abbreviated as follows: Guiyang in 2019 (GY19), Guiyang in 2020 (GY20), Zhucheng in 2019 (ZC19) and Zhucheng in 2020 (ZC20).

**Fig. 1.**
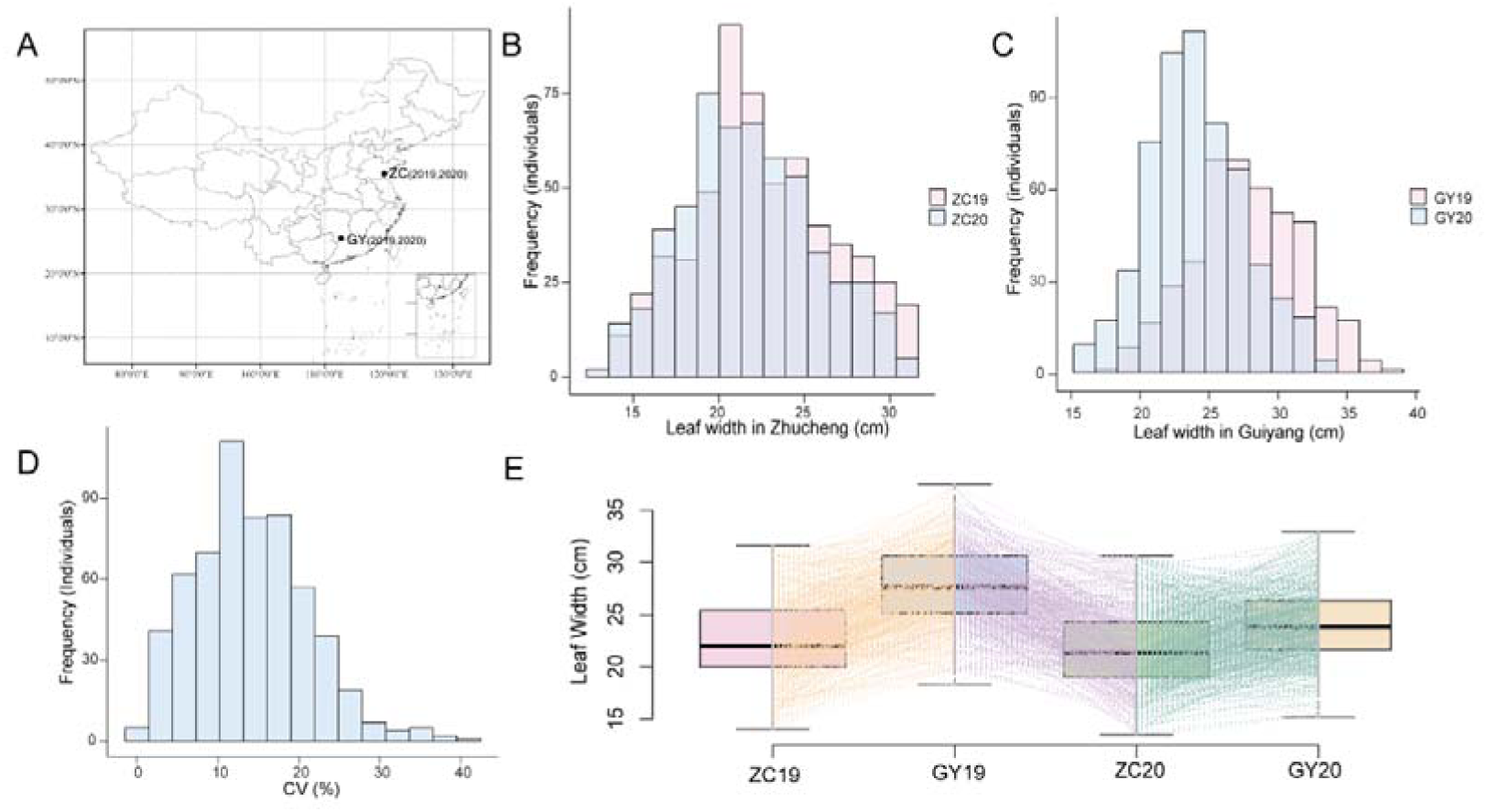
Variation in leaf width of MAGIC population across environments. **A)** Geographical illustration of two filed plantations, Zhucheng and Guiyang, where the MAGIC population was phenotyped. **B)** Leaf width phenotype distribution histogram of ZC19 and ZC20. **C)** Leaf width phenotype distribution histogram of GY19 and GY20. **D)** Distribution histogram of CV of phenotype. **E)** Boxplot of LWm measured at four environments.

Phenotypic analysis across different environments indicated that tobacco leaf width exhibited continuous variation, ranging from 13.5 cm to 37.5 cm. Environmental changes significantly altered the mean across the four environments, with a widespread reaction norm observed for LWm. For example, when the environment shifted from ZC20 to GY19, the leaf width in Guiyang (LWm = 27.8 cm, CV= 0.14, Fig. 1D) showed a marked increase compared to Zhucheng (LWm = 21.8 cm,CV = 0.15), resulting in variation in LWp (Fig. 1E).

When location was held constant, the leaf width in GY20 averaged 24.1 cm, representing a 13% reduction compared to GY19. Similarly, in ZC20, the LWm reached at 21.8 cm, reflecting a 4% decrease relative to ZC19. Within the same year, the LWm in GY19 was 27.8 cm, 23% greater than that in ZC19, whereas in 2020, the LWm in GY20 reached at 24.1 cm, 11% higher than in ZC20. Overall, the variation in LWm across different locations (up to 23%) was substantially greater than the interannual variation at the same location (4%–13%), indicating that geographic location had a more pronounced effect on LWm (Fig. 1B, C). Nevertheless, year-to-year fluctuations were also evident, suggesting that LWm is highly responsive to environmental changes and exhibits significant phenotypic plasticity.These results indicate that environmental changes affect both LWm and LWp.

To further dissect the phenotypic variance components, we performed analysis of variance (ANOVA) considering genotype (G), environment (E), and genotype-by-environment (G×E) interactions. The results revealed that genotype explained 40.27% of the total variance in leaf width, while environment contributed 12.02%. Notably, the G×E interaction accounted for 31.29%, indicating a significant role of environmental responsiveness in shaping leaf width variation (Fig. S1). These findings suggest that genetic effects predominantly determine leaf width, but substantial G×E interactions underscore the importance of plastic responses to environmental fluctuations.

### 3.2 Environmental perturbation buffers/releases additive variation for LWm by altering the underlying genetic architecture

Moderate variation (0.13–0.61; Fig. 2A, Fig.S2) in the estimated narrow-sense heritability (*h^2^*) across different environmental conditions was observed. Since phenotypic variation (Vp) varied significantly among the four environments (Fig S3), the amount of additive genetic variation (Va), estimated as Vp × *h^2^*, ranged from 4.1 to 11.2 (Fig. 2B). These fluctuations in Va were likely driven by changes genetic effects from one or more QTLs or by the activation and deactivation of a completely new set of QTLs, indicating that the genetic architecture of LWm was dynamic across the four environments.

**Fig. 2.**
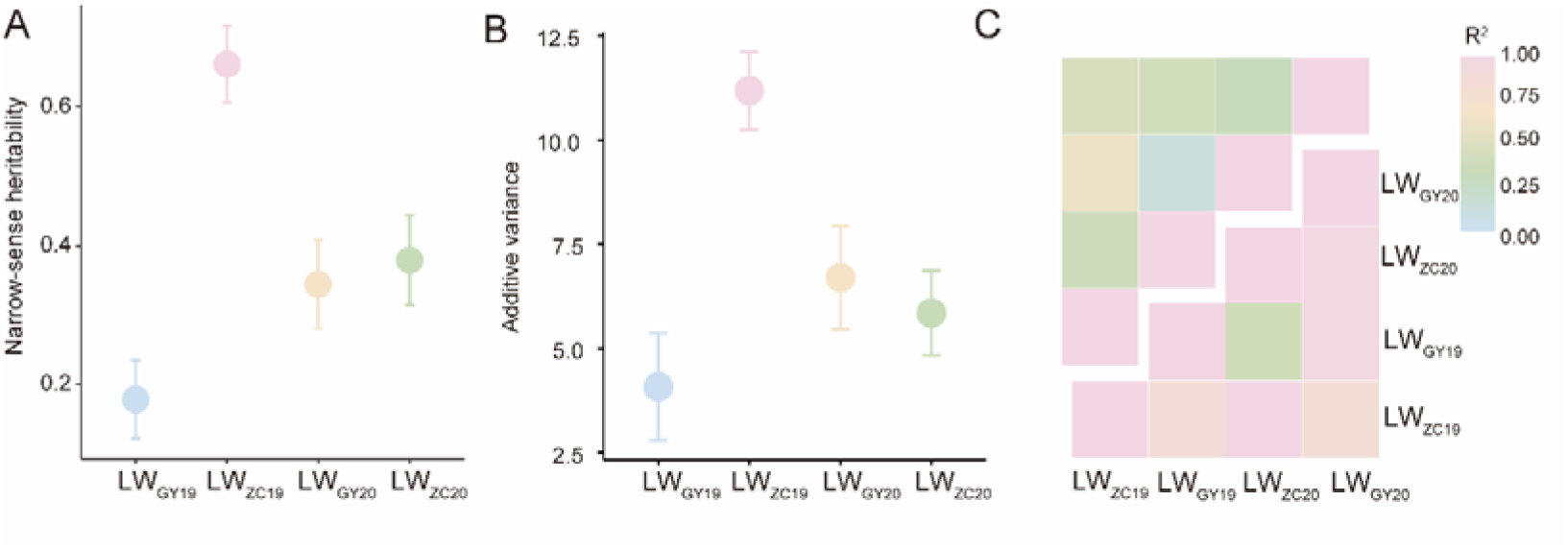
Variation of genetic effects across four environments. **(A)** Estimated narrow-sense heritability (*h^2^*) for LWm at each environment. Each dot represents a point estimate of *h^2^* indicated on the y-axis and the corresponding environment was labelled in the x-axis. Standard errors were plotted as error bars. (**B)** Estimated additive variance at each environment. (**C)** Upper triangular: Pairwise Spearman rank correlation among LWm measured under four environments. (**C)** Lower triangular: Pairwise genetic correlation among LWm measured under four environments.

This hypothesis was supported by the variation in pairwise genetic correlations from the multivariate mixed model analysis (Fig. 2C). The estimated genetic correlation between environments was as low as 0.4, suggesting considerable changes in polygenic scores and breeding values across individuals. These changes may be attributable to genotype-environment interactions, where certain alleles or QTLs exert different effects in different environments. Collectively, these results illustrate that the genetic architecture of LWm varied across these environments due to environmental changes. In the following sections, we will quantify this variation and dissect the genetic basis underlying LWm variation across environments.

### 3.3 Polygenic architecture underlying tobacco LWm variation and LWp variation

A widespread reaction norm was observed for LWm. At the population level, the median LWm in Zhucheng was significantly smaller than that in Guiyang (Fig. 1B), resulting in variation in LWp. Therefore, we quantified LWp using two types of metrics. The first type included two measures: Finlay–Wilkinson Regression (FWR), which evaluates genotype-by-environment interactions, and the across-environment variance in rank-transformed LWm (VarR), which quantifies the overall plasticity (OP of LW across all environments. The second metric involved assessing pairwise LWm differences for all possible pairs of environments, quantifying specific changes across paired environments, resulting in a total of six specific plasticity (SP) measures.

Almost all plasticity measurements showed a continuous distribution with low to high narrow-sense heritability (0.015–0.61, Fig. S2, Fig. S4). These results indicate that LWp is a heritable trait influenced by both genetic and environmental factors, providing a solid foundation for further dissection of its genetic architectureA total of 45 QTL were identified for LWm and LWp by performing GWAS (Fig. 3A, Table S2, Fig S5, Fig S6), including 14 QTL for LWm, 9 QTL for lead width BLUP (LW_BLUP_) and 43 QTL for LWp, LWm and SP share 12 overlapping loci, accounting for 27% of all identified QTLs. Additionally, LWm and BLUP exhibit complete overlap at 9 loci. Furthermore, LWm, SP, and BLUP share 8 common loci, representing 18% of all QTLs. These results indicate a high degree of overlap among the three methods, indicating a shared genetic basis between LWm, LWp and BLUP (Fig. 3C, Table S3). The majority of loci show a high degree of overlap across different methods. This may be due to the variation in the strength of genotype-environment interactions in the environment, which leads to partial differences in the detected QTLs and ultimately to the phenotypic plasticity of these traits. By contrasting genetic effects of QTL across sites, two types of QTL, whose effects changed in magnitude, were associated with LWp For example, *qLW1-3* showed a significant effect for LW_ZC19_ (5.6 ± 0.7cm; *P* = 4.8 × 10^-16^; Fig. 3D), LW_ZC20_ (3.7 ± 0.6cm; *P* = 4.4 × 10^-9^) as well as on the specific plasticity measures LW_GY19-ZC20_ (*P* = 5.9 × 10^-8^; Fig. 3D) and LW_ZC19-GY19_ (*P* = 2.2 × 10^-14^; Fig. 3D), indicating changes in the magnitude of genetic effects contributed to the variation of LW plasticity. In contrast, *qLW9-1* (Fig. 3E), was exclusively detected for several LWp but was not associated with any LWm or LW_BLUP_ measurements. Altogether, these results suggested that changes in magnitude of genetic effects across sites caused variation in plasticity. The changing genetic effects highlighted the role of QTL by environment interaction in the variation of LWm and LWp.

**Fig. 3.**
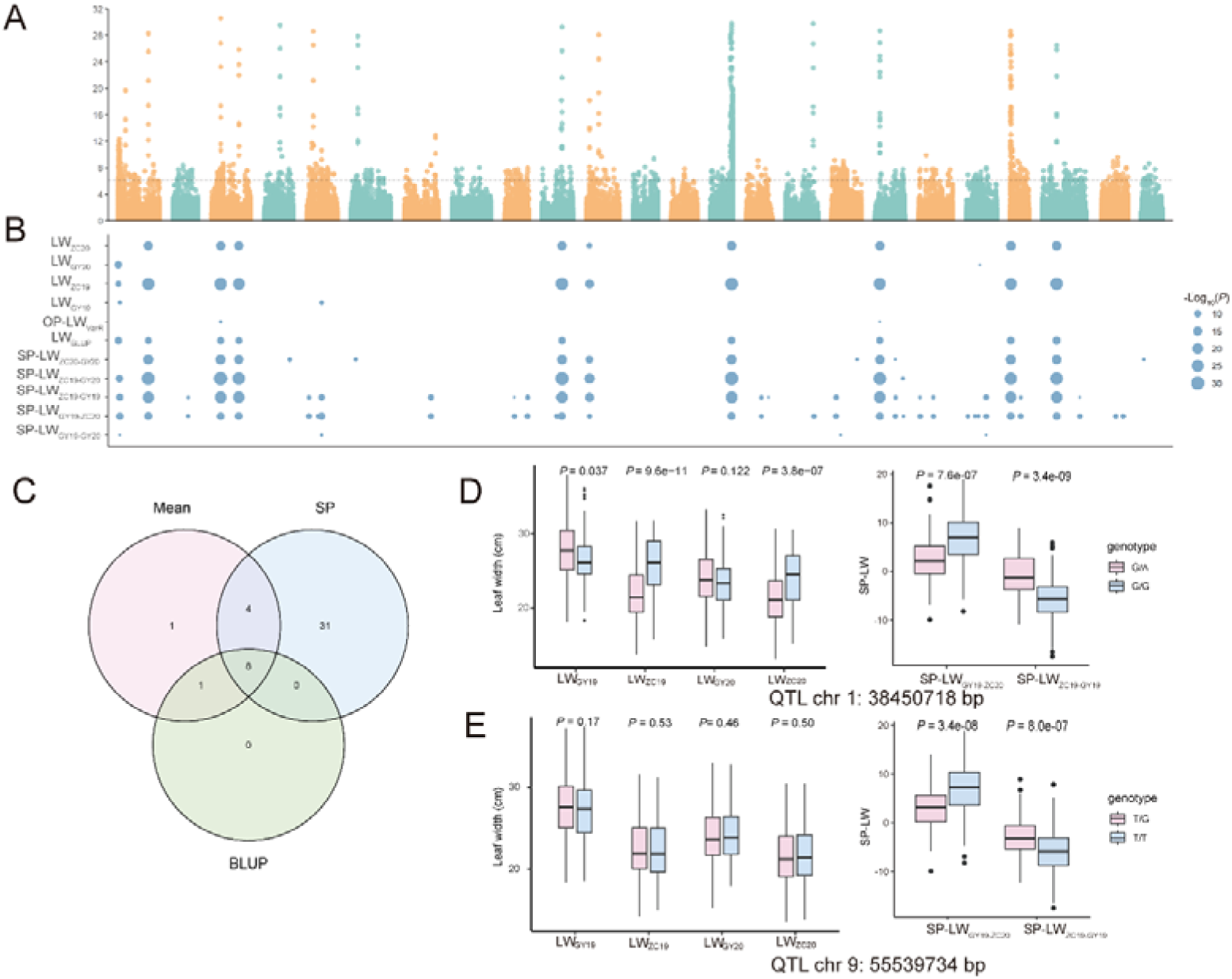
Summary of the QTLs associated with mean and plasticity measures for LW. **(A)** Manhattan plots overlaying genome-wide association analysis results for the mean, plasticity measurements and BLUP for LW. The black horizontal dashed line indicated the Bonferroni-corrected genome-wide significance threshold derived as 0.05/Me (Me is the effective number of independent markers; Methods) **(B)** QTL associated with the LWm and LWp measured at four environments and the LW_BLUP_ (y-axis). Each dot represents a SNP, and the size of the dot is proportional to its - Log10 p value, as indicated in the legend on the right. Loci with a p value above the genome-wide significance threshold are colored in blue. **(C)** Venn diagram illustrating the overlap of QTL detected for the 3 types of LW measurements. **(D) and (E)** Genotype-to-phenotype maps for LWm and LW_SP_ at Chr 1: 38450718bp, Chr 9: 55539734bp.

### 3.4 qLW14 associated with LWm and LWp measurements

We detected a QTL, *qLW14* located at chromosome 14: 116-123 Mb, associated with LWm at Zhucheng (LW_ZC19_, LW_ZC20_) and multiple LWp measurements (SP-LW^GY19-ZC20^, SP-LW^ZC19-GY19^, SP-LW^ZC19-GY20^, SP-LW^ZC20-GY20^, Fig. 4A, Fig S5). Linkage disequilibrium (LD) analysis revealed a high level of LD in this region (Fig. 4B). Within the high level LD block, we identified three major haplotypes (hap1-3). Phenotypic comparisons of these haplotypes under two environmental conditions showed that in Zhucheng, lines carrying hap2 exhibited approximately 16% wider leaves compared to hap1 and hap3 (Fig. 4C), whereas no significant differences were observed in Guiyang (Fig. 4D). Further analysis revealed that the hap2 haplotype originated from the parent *Vam* which harbored a ∼3 Mb deletion between 121 and 124 Mb, overlapped with the *qLW14* region (Fig. 4E).

**Fig. 4.**
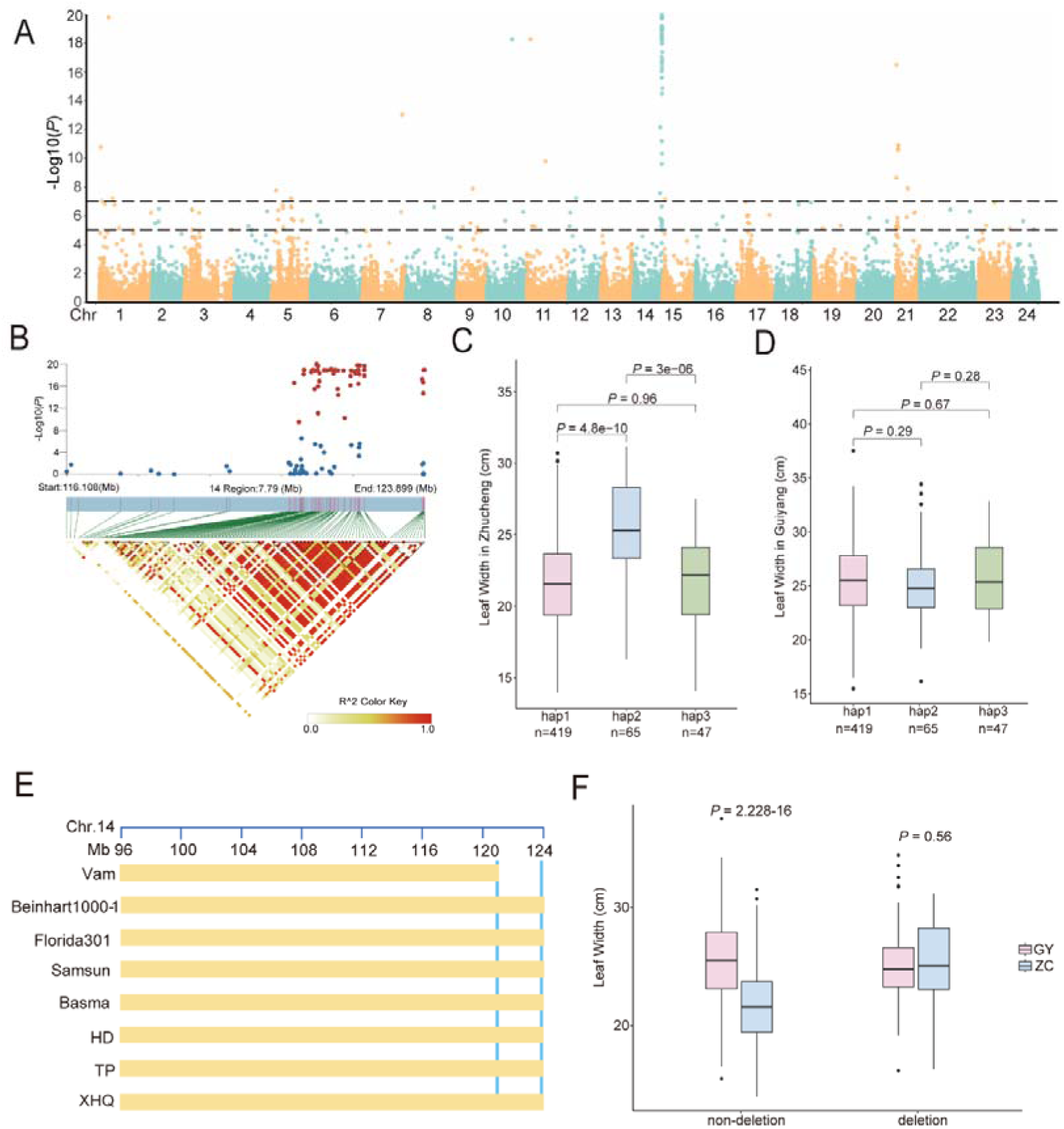
GWAS of the LW_ZC19-GY20._ **(A)** Manhattan plot for plasticity measurement LW_ZC19-GY20_. The black horizontal dashed line represents the Bonferroni-corrected genome-wide significance threshold of 0.05/Me (Me is the effective number of independent markers; Methods). **(B)** LD heatmap showing the region surrounding a strong peak (chromosome 14: 116 Mb–124 Mb). One-way ANOVA was used to compare the leaf width of three haplotypes lines *qLW14* in the year of 2019 and 2020 at Zhucheng, Shandong **(C)** and Guiyang, Hunan **(D).(E)** Haplotype analysis from re-sequenced eight parents for MAGIC population (*Vam*, *Beinhart1000-1*, *Florida301*, *Samsun*, *Basma*, HD: *honghuadajianyuan*, TP: *tangpeng*, XHQ: *xiaohuaqing*) revealed a 3Mb deletion in *Vam*. Difference in leaf width of the deletion and non-deletion*qLW14* in the year of 2019 and 2020 at Zhucheng, Shandong and Guiyang, Hunan **(F).**

To explore the phenotypic effect of this deletion, we compared leaf width between deletion and non-deletion lines in both environments. Phenotypic analysis showed that, in Guiyang, lines without the deletion exhibited leaves approximately 18.5% wider than those grown in Zhucheng. In contrast, delete that fragment did not significantly affect leaf width in either environment (Fig. 4F). Further analysis revealed that 125 genes were annotated within this region, including 15 tandemly duplicated genes encoding MADS-box proteins. Members of the MADS-box family have been reported to regulate plant development in response to light and temperature. Overall these results highlighted a general contribution from *qLW14* by environment interaction to variation of LWp.

### 3.5 Accounting for dynamics in genetic architecture improved LWm prediction across environment

To improve the effectiveness of cross-environment prediction of leaf width traits, we evaluated the potential of integrating genetic diversity, environmental variation, and their interaction in complex trait prediction by jointly modelling genotype, environment, and their interaction (referred to as the GEAI model, Methods). First, we explored the ability to predict untested lines (Newly developed lines) at the four environments using five-fold cross-validation. Compared to GBLUP, which provides a universal prediction for all sites, our model not only offered site-specific predictions but also improved prediction accuracy. The average prediction accuracy for LW increased by 8.7%. (Fig. 5A, 5B).

**Fig. 5.**
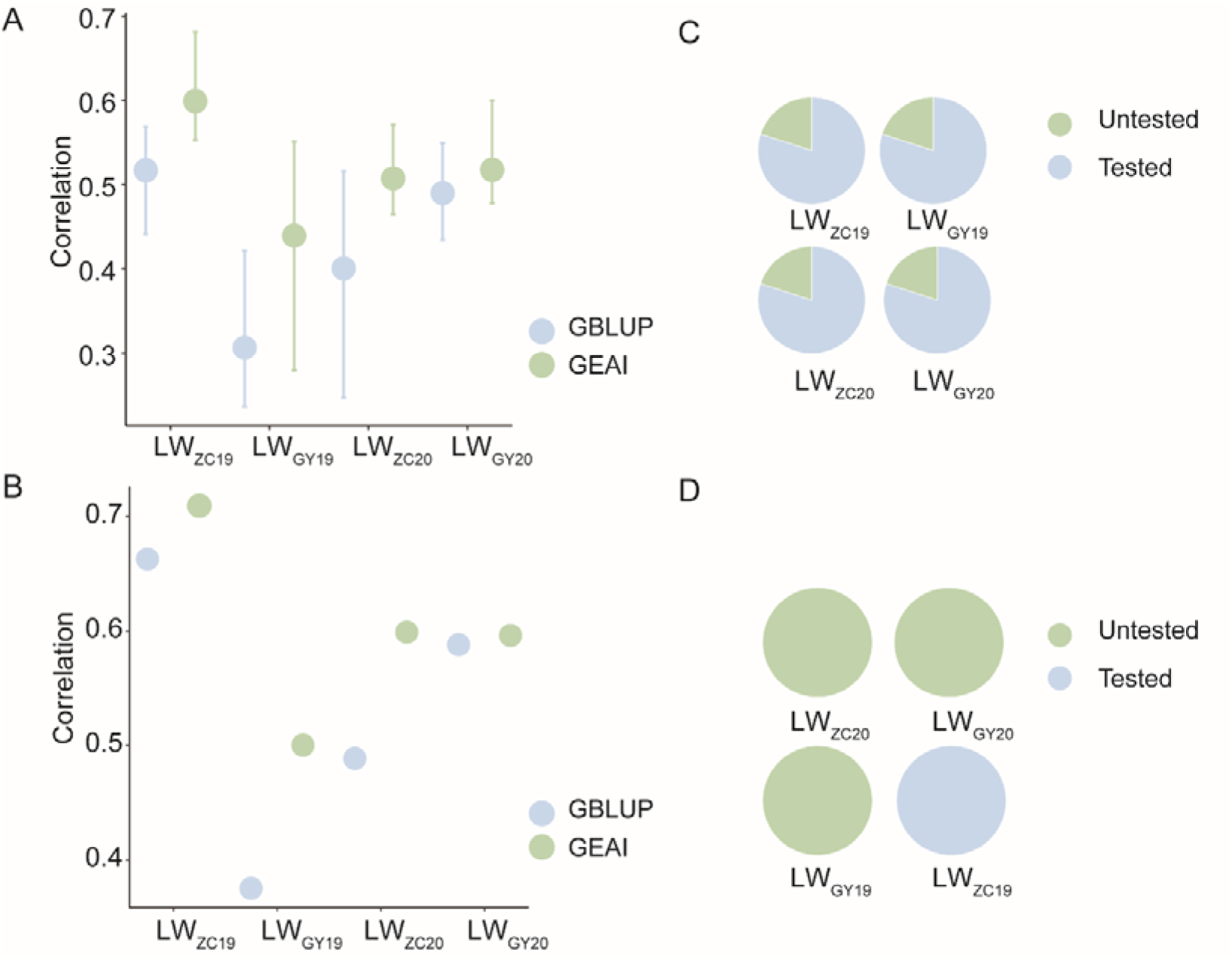
Prediction of leaf width phenotype using GEAI model. **(A)** Use 5-fold cross-validation to predict the phenotypes of the remaining 20% lines in each environment. **(B)** Use the phenotypes of all strains in three environments in the four environments as training set for LW, and the remaining environments as a test set. **(C)** Schematic illustration of the prediction design for **(A)**. **(D)** Schematic illustration of the prediction design for **(B)**, with LW_ZC20_, LW_GY20_, and LW_GY19_ as training set and LW_ZC19_ as the tested set.

In the second scenario, we performed prediction for novel sites without any tested lines by utilizing phenotypic data from three out of the four environments as the training set, while the phenotype data from the remaining environment served as the test set. Compared to the GBLUP model, our GEAI model exhibited an average improvement of 7.2% in prediction accuracy (Fig. 5C, 5D). This substantial enhancement underscores the effectiveness of our GEAI model in capturing and leveraging genotype-environment interactions to enhance prediction accuracy across different environmental contexts. Overall, our study highlighted the potential of integrating QTL by environment interaction in understanding complex traits variation and predictions.

## 4. Discussion

### 4.1 Genetic and Environmental Regulation of Leaf Width Plasticity

Consistent with studies in maize and rice (Guo et al., 2020; Ji et al., 2023), in this study, we found that 33% of the QTLs associated with LWp were shared with those associated with LWm. This overlap suggests a common genetic basis between LWm and LWp, indicating that some genetic mechanisms controlling leaf width are conserved across different environmental conditions. These shared QTLs may regulate fundamental developmental processes that influence leaf morphology. However, despite this genetic overlap, the observed phenotypic differences between LWm and LWp emphasize the important role of phenotypic plasticity in allowing plants to adapt to varying environmental conditions.

Due to the lack of detailed environmental factor data in our study, we were unable to definitively determine which specific environmental factors, such as soil moisture, temperature, light intensity, or nutrient availability contribute to the plasticity of leaf width. Environmental factors often interact in complex ways to shape phenotypic outcomes, and their effects may vary depending on the plant’s developmental stage or the intensity and duration of stress. For example, drought stress may trigger specific signaling pathways that alter leaf development, while temperature fluctuations could influence cell expansion and division rates, further modulating leaf width. Additionally, epigenetic modifications, such as DNA methylation or histone modifications, could mediate the plant’s response to environmental cues, potentially influencing phenotypic plasticity (Guo et al., 2025; Liu et al., 2022a; Ma et al., 2024; Prasad et al., 2015; Yang et al., 2025). Further research is needed to explore the specific roles of environmental factors in the variation of LWm and LWp.

### 4.2 Potential role of MADS-box transcription factors at the qLW14 locus in leaf development

Haplotype analysis of the *qLW14* locus revealed the presence of multiple duplicated MADS-box transcription factors. The MADS-box gene family has been shown to play a role in leaf development in both *Arabidopsis* and *tomato*. In *Arabidopsis*, the MADS-box member FRUITFULL (FUL) regulates leaf morphology by modulating vascular bundle development (Bemer et al., 2017), while in *tomato*, SlMBP21 influences leaf development by suppressing genes associated with seed development (Wang et al., 2021). These findings suggest that MADS-box genes may have conserved functions in leaf development.

Although the candidate interval of *qLW14* remains relatively large, advancements in multi-gene editing technologies, such as CRISPR/LbCpf1 (Li et al., 2020), now make it feasible to perform fine mapping and functional validation of potential candidate genes within this QTL. Future research could integrate gene editing, expression analysis, and genetic mapping to narrow down the candidate region and identify key genes, thereby elucidating the molecular mechanisms through which MADS-box genes regulate leaf development. These studies will not only enhance our understanding of the genetic regulatory networks underlying leaf development but also provide valuable genetic resources for crop breeding.

### 4.3 Optimization of Genomic Prediction Model Incorporating Genotype-Environment Interactions

We developed a genomic prediction model that incorporates genotype-environment interactions. When using data from three environments as the training set to predict phenotypes in the remaining environment, this model outperformed the traditional GBLUP. This superiority is mainly due to the model’s ability to capture interaction effects between detectable QTL and environmental conditions. However, due to the lack of diverse environmental factor data, we were unable to fully elucidate the specific interaction mechanisms between QTLs and particular environmental factors, which warrants further investigation.

Despite its limitations, we recommend prioritizing the GEAI model for cross-environment trait prediction, as it significantly enhances prediction accuracy. By establishing a prediction model that is responsive to environmental changes, we can effectively predict phenotypes, such as leaf width, across different environments. This approach lays the foundation for improving the accuracy of genome-wide predictions of agronomic traits across multiple environments. Future research could integrate more environmental factor data (e.g., temperature, humidity, light) to further optimize model parameters and explore the specific interaction mechanisms between QTLs and environmental factors. Such efforts will provide essential theoretical support and technical tools for precise phenotype prediction and genotype selection in crop breeding.

## Author Contributions

Conceptualization, L.L., W.L., Y.H., R.H.; methodology, Y.H., R.H., S.Y., J.Y., G.J., Y.Z., H.S.; formal analysis, L.L., W.L., Y.H., W.D.; investigation, Y.S., L.X., Z.W., W.Z., L.C.; data collection and curation, L.C., S.Z., Y.Z., H.S., W.Z., Z.J., Y.S., L.G.; writing original draft preparation, L.L., W.L.; writing reviewing and editing, L.L., W.L., Y.Z., Z.C., H.S.; visualization, L.L., W.L, Y.H; supervision, H.H., Y.Z., H.S.; funding acquisition, Y.Z., H.S., L.C.

All authors have read and agreed to the published current version of the manuscript.

